# Structural basis for transcription activation through cooperative recruitment of MntR

**DOI:** 10.1101/2024.06.28.601288

**Authors:** Haoyuan Shi, Yu Fu, Vilmante Kodyte, Amelie Andreas, Ankita J. Sachla, Keiki Miller, Ritu Shrestha, John D. Helmann, Arthur Glasfeld, Shivani Ahuja

**Author notes:** Coauthors with equal contributions (Vilmante Kodyte, Amelie Andreas).

## Abstract

The manganese transport regulator (MntR) from *B. subtilis* is a dual regulatory protein that responds to heightened Mn^2+^ availability in the cell by both repressing the expression of uptake transporters and activating the expression of efflux proteins. Recent work indicates that, in its role as an activator, MntR binds several sites upstream of the genes encoding Mn^2+^ exporters, leading to a cooperative response to manganese. Here, we use cryo-EM to explore the molecular basis of gene activation by MntR and report a structure of four MntR dimers bound to four 18-base pair sites across an 84-base pair regulatory region of the *mneP* promoter. Our structures, along with solution studies including mass photometry and *in vivo* transcription assays, reveal that MntR dimers employ polar and non-polar contacts to bind cooperatively to an array of low-affinity DNA-binding sites. These results reveal the molecular basis for cooperativity in the activation of manganese efflux.

## Introduction

Transition metals are essential to the function of bacterial cells but become toxic when overabundant. That dual character requires careful monitoring of transition metal availability in the cytoplasm, a function generally performed by metal-responsive regulatory proteins^1–3^. For some metal ions, such as Ni^2+^ and Zn^2+^, a bacterium may possess coupled regulatory proteins. For example, at low zinc availability in *E. coli*, uptake is derepressed by Zur, but at high zinc availability, zinc efflux is activated by ZntR^4^. Similarly, NikR negatively regulates Ni^2+^ uptake in response to nickel, while RcnR is induced by higher concentrations of Ni^2+^ to promote Ni^2+^ export^5^. These coupled systems act in concert to tightly maintain metal ion homeostasis in the cell. In the instance of iron, the metalloregulators Fur and DtxR are capable of both repression and activation of gene expression under elevated iron availability in the cell^6,7^. In the case of repression, it is understood that Fur or DtxR binding to an operator limits RNA polymerase access to a promoter. It is unclear, however, if activation is achieved by recruiting RNA polymerase to a promoter adjacent to the operator, or whether it is achieved by coordination with other regulatory systems.

Recently, it was discovered that manganese homeostasis in *B. subtilis* can also be maintained by repression and activation through the action of a single regulatory protein, MntR^8^. MntR, a distant homolog of DtxR, is a dimer of 142-residue subunits that forms a binuclear complex with two Mn^2+^ ions per subunit stabilizing a high-affinity DNA-binding conformation of the protein^9^. At high manganese availability in *B. subtilis* cells, the expression of two uptake transporters, MntH, a proton-coupled NRAMP family transporter, and MntABCD, an ATP-binding cassette transporter, is repressed. As manganese levels rise further, the expression of MneP and MneS, primary and secondary Mn^2+^ efflux transporters, is activated^8^. Repression by MntR has been extensively studied. Under conditions of Mn(II) sufficiency (∼10 µM) a single MntR dimer binds to duplex DNA containing either the *mntA* or *mntH* operator with nanomolar affinity^10^. Those operators overlap the ***σ***^A^-dependent promoter upstream of the respective open reading frames, and thus MntR can repress expression by acting as a molecular doorstop^10^.

Activation of transcription by MntR is a more recent discovery and the molecular basis of activation is not well understood. Previous *in vivo* results^8^ suggested cooperative binding of multiple MntR dimers to three sites across ∼80 base pairs on both the *mneP* and *mneS* promoters is necessary for activation of transcription. For both *mneP* and *mneS,* one of the predicted MntR binding sites overlaps the *σ*^A^ promoter -35 consensus site for RNA polymerase (RNAP) binding.

In this study, we present two single-particle cryogenic electron microscopy (cryo-EM) structures of MntR bound to the regulatory region of the *mneP* operon. Our structures reveal the stoichiometry of MntR-DNA binding and the structural basis of cooperative binding under conditions of manganese sufficiency. Through structural, biochemical, and in vivo expression studies, we identified crucial protein-protein interactions required for the cooperative recruitment of MntR dimers to the *mneP* promoter and necessary for gene activation.

## Results and Discussion

### Four MntR dimers bind the regulatory region of the efflux *mneP* promoter

We prepared a sample for cryo-EM (detailed in the methods), with an 84-bp DNA duplex containing the *mneP* regulatory region (referred to as P84; Supplementary Table 1, Supplementary Fig. 1) with a 4-fold excess of MntR dimers in the presence of 1 mM Mn^2+^ ions. From a single cryo-EM dataset obtained from this sample, we obtained two separate three-dimensional (3D) reconstructions (Supplementary Fig. 2). At first the data processing was limited to a smaller image extraction box size of 255 Å, resulting in a 3.09 Å reconstruction featuring well-resolved density for two MntR dimers bound to a portion of the P84 duplex, which we refer to as the (MntR_2_)_2_-P84 map (Supplementary Fig. 3a, Supplementary Table 2). Further analysis of the cryo-EM micrographs, using the (MntR_2_)_2_-P84 map as a template and extracting at a larger box size of 852 Å, yielded a second 3D reconstruction at a nominal resolution of 4.17 Å. Instead of the expected three occupied binding sites, the refined map shows density for four MntR dimers bound to P84, which we named (MntR_2_)_4_-P84 map (Supplementary Fig. 3b, Supplementary Table 2). While the flexibility of the DNA induced heterogeneity and lowered resolution, the (MntR_2_)_4_-P84 map nevertheless provides evidence towards the stoichiometry of MntR homodimers to the full regulatory region of the *mneP* promoter sequence. Notably, this complex is consistent with the three MntR-binding sites previously shown to be essential for *in vivo* activation of *mneP* and suggests that they function together with a fourth site^8^.

#### Model refinement, metal ions, and DNA register

The refined structures of MntR dimers bound to DNA are very similar to the structure of MntR crystallized in the absence of DNA^11^ (PDB: 2F5C; Supplementary Fig. 4). Structural alignment to the two MntR dimers fit to the (MntR_2_)_2_-P84 complex map to the starting MntR model (PDB: 2F5C) yields RMSDs of 0.8 Å and 0.9 Å. When the flexible wing, which is in a slightly different position when bound to DNA, is excluded from alignment, the RMSDs are reduced slightly to 0.7 and 0.8 Å. The distance between the N-terminal DNA-binding domains can be measured by the separation of dyad-related alpha-carbons of Lys41 in the recognition helix. The separation in the starting model (PDB: 2F5C) is 32.1 Å and in the two dimers fit to the (MntR_2_)_2_-P84 map, 32.5 Å and 33.5 Å (Supplementary Fig. 4).

A binuclear manganese binding site has been observed for all manganese complexes of MntR and its homologs both in crystal structures and in solution^1,11–14^, which has been hypothesized to be essential for full activation of MntR for DNA binding^15^. In our cryo-EM maps, good density is generally present for the A-site metal, especially in the more central subunits of each complex, but is somewhat equivocal for the C-site metal. Nevertheless, both metals were modeled in all subunits, and geometry restraints, based on an existing structure of MntR bound to Mn^2+^ (PDB: 2F5D)^11^, were used in refinement.

Modeling the 84-bp duplex containing the *mneP* operator region was complicated by its asymmetry. DNA orientation in individual particles can be in either direction, and the end bases are not present in either map. In each model, we have chosen to model one orientation of the DNA duplex. In the (MntR_2_)_4_-P84 model, the duplex contains base pairs 3/-3 through 79/-79. In the (MntR_2_)_2_-P84 model, the DNA duplex is modeled using base pairs 23/-23 (along with an overhanging base at position 22) through 60/-60, representing, roughly, the central 38 base pairs of the complex. That choice acknowledges not only the orientation issue described above but also that the particles chosen to generate the (MntR_2_)_2_-P84 map may come from either the center or ends of the full complex.

A further consideration in modeling the DNA, as described above, is the specific register for the positions of the MntR dimers on the sequence. Previous work^8^ identified three individual MntR binding sites in both the *mneP* and *mneS* operators. The (MntR_2_)_4_-P84 map and mass photometry (MP) solution data (see below) clearly indicate that MntR dimers bind to four sites across the P84 sequence. Accordingly, we have re-evaluated the operator sequences and have identified a fourth sequence showing similarity to the sites previously identified (Supplementary Fig. 1). In our revised analysis of the *mneP* and *mneS* regulatory regions, we assign the sites from 1-4, with site 1 overlapping the RNA polymerase (RNAP) binding site, and site 4 most distant from the promoter (Fig. 1a). The position of MntR dimers on duplex DNA is consistent with each MntR interacting with nucleic acid bases across 18 bp, with 9 bp inverted repeats. These 18-bp binding sites are each separated by 1 bp. The protein-DNA duplexes have been modeled such that each MntR subunit is situated equivalently to its corresponding half-site, and the 2-fold axis relating to each MntR dimer is aligned with the 2-fold axis relating the two 9-bp inverted repeats in each operator binding site (Fig. 1a, b).

**Figure 1.**
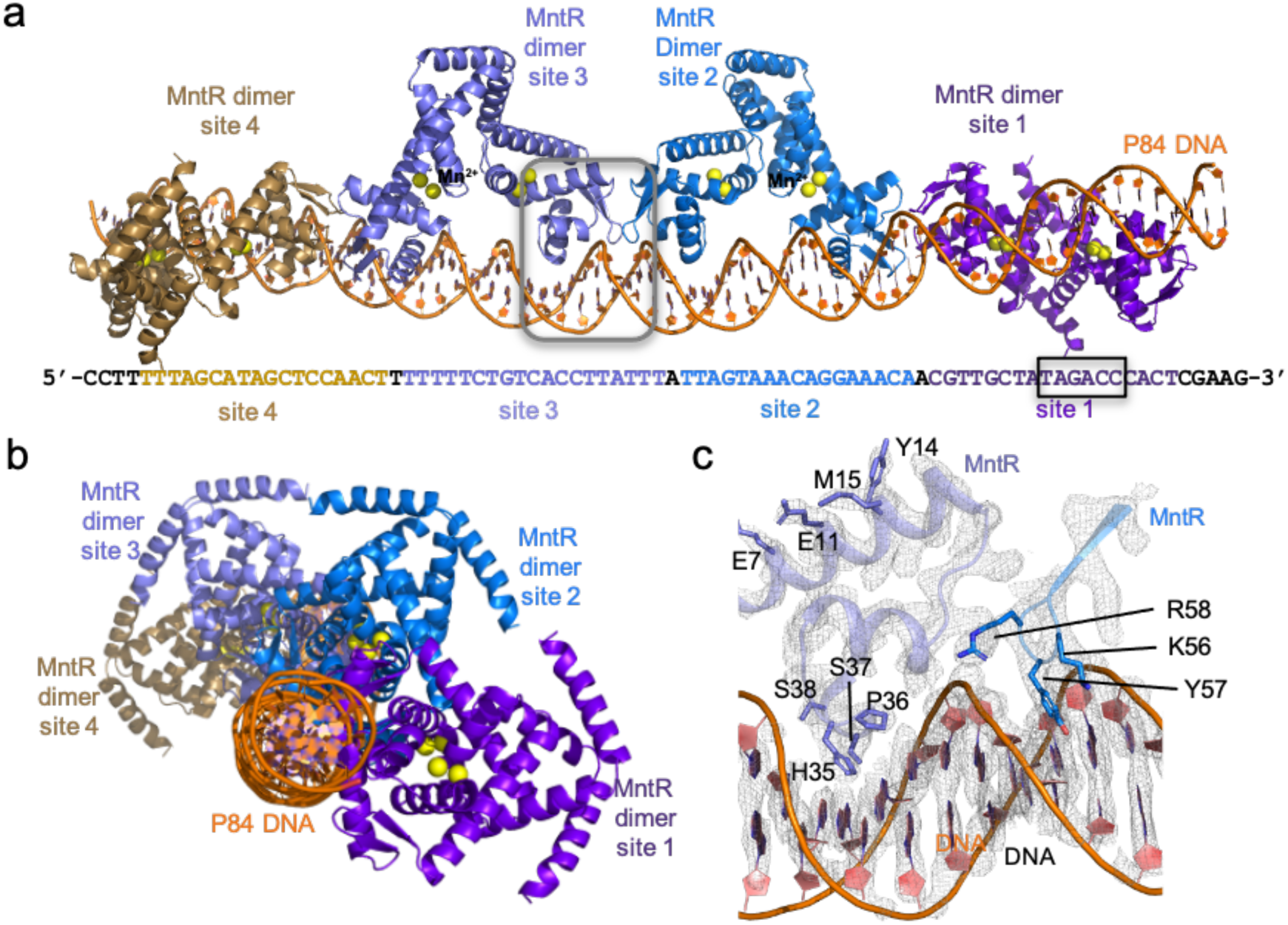
Structure of (MntR_2_)_4_-P84 complex. (a) Cryo-EM structure of (MntR_2_)_4_-P84 complex showcasing the organization of the four MntR dimers with respect to P84. The P84 sequence is displayed below highlighting the four 18-bp MntR dimer binding sites (site 1-4) in color. The black box highlights the RNA polymerase (RNAP) binding site (-35). (b) An alternate view of the (MntR_2_)_4_-P84 structure after a 90° rotation compared to (a) looking down the DNA helical axis. (c) A close-up view of the boxed region from (a) showcasing the electron density for some of the amino acids in the MntR dimer-dimer and MntR-DNA interface.

#### DNA conformation in the (MntR_2_)_4_-P84 complex

Generally, the DNA duplex adopts the B-conformation in both models with small deviations from overall linearity. Given the constraints applied during refinement and the moderate to low-resolution maps, no detailed analysis of DNA conformation is warranted. However, there is a distinct, but light bending of the duplex visible in both models, but more pronounced throughout the 77 base pairs included in the (MntR_2_)_4_-P84 model. *Curves+* software^16^ was used to analyze the duplex. When comparing the best-fit linear axis for the DNA double helix (Supplementary Fig. 5) to a curve that traces the helical axis as it alters its path with each base pair, the gentle curvature of the duplex is evident. The DNA bends slightly towards the bound MntR dimers, with slight inflection points at the dimer-dimer interface points, suggesting that alterations in DNA conformation assist in the formation of contacts between MntR dimers.

### Protein-DNA interactions include major and minor groove contacts

The cryo-EM structures of the MntR-DNA complexes presented here provide a starting point for defining protein-DNA interactions that promote specific recruitment of MntR to its operators. As noted above, each MntR dimer binds to an 18-bp site consisting of a pair of 9-bp inverted repeats with an average buried surface area of ∼1460 Å^2^ for each MntR dimer at the protein-DNA interface (Fig. 2). Overall, each dimer spans 20-bp through contacts made by residues of the wing motif (residues 52-62) to the phosphodeoxyribose backbone (Supplementary Fig. 4). Comparison of sites 1-4 of the *mneP* operator region with the comparable expected sites for *mneS* as well as for the *mntA* and *mntH* operator regions permits a sequence alignment that highlights preferred base pairs within the half-site recognized by MntR (Supplementary Fig. 1). That alignment then permits interpretation of the protein-DNA interactions that can be inferred from the (MntR_2_)_2_-DNA complex structure, which offers a < 3.0 Å resolution between the central two MntR protomers (chain F and G) and the DNA duplex (Fig. 1c, Supplementary Fig. 3). Although modeling of the DNA sequence is complicated by the lack of symmetry in the DNA duplex used in cryo-EM studies, the positions of residue side chains relative to base pairs indicate which residues can participate in selective binding, while non-sequence specific interactions to phosphates can be defined. Also, the A_7_T_-7_ base pair is conserved in all half-sites in the DNA duplex used in this study and can be more confidently modeled (Fig. 2 and Supplementary Fig. 1b).

**Figure 2.**
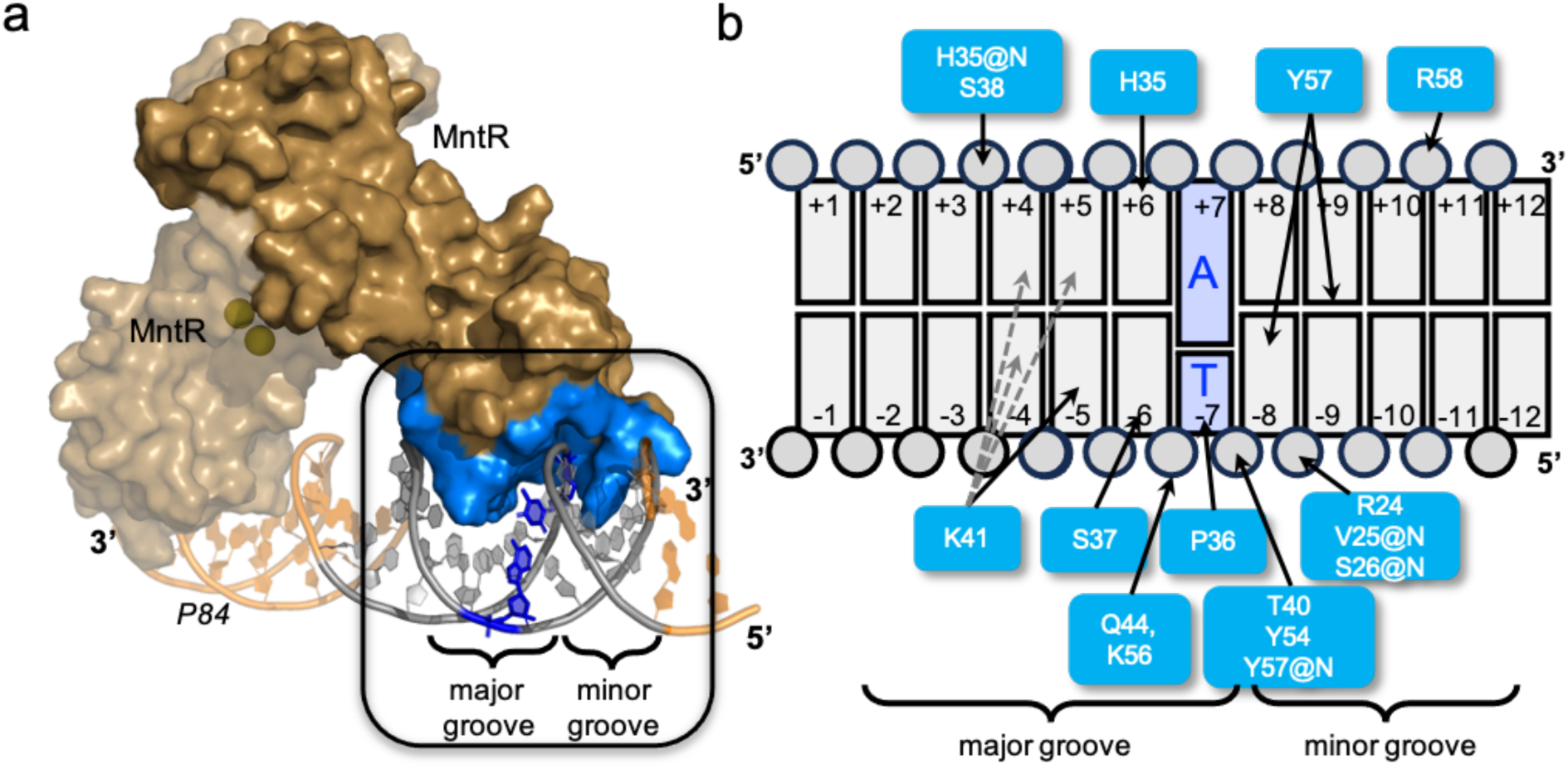
MntR-DNA interactions. (a) A portion of the cryoEM structure of the (MntR_2_)_2_-P84 complex highlighting a single dimer of MntR in surface representation (tan) bound to an 18-bp operator region on P84. Mn^2+^ ions are presented as yellow spheres. The buried surface area of an MntR monomer is highlighted in blue interacting with a major and minor groove of P84 highlighted in grey. (b) A schematic presenting the interactions between amino acids of MntR (blue) with the nucleotides of P84 (grey) in the boxed region in (a). The base pair at -7/+7 position is always a TA base pair (highlighted in dark blue in (a) and (b)) in all 8 half sites in the (MntR_2_)_4_-P84. The label “@N” indicates an interaction with the backbone amide nitrogen. His35, Pro36, Ser37 and Lys41 interact with the base edges in the major groove and Tyr57 interacts within the minor groove.

The MntR recognition helix (residues His35-Lys48) lies within the major groove of DNA, placing the side chains of residues His35, Pro36, Ser37, and Lys41 in proximity to the edges of base pairs 4/-4 through 7/-7 in both half-sites (Fig. 2 and Supplementary Fig. 4). Lys41 on subunit G is positioned to donate a hydrogen bond to the base at position -5, while the map is more ambiguous for Lys41 on the F subunit, suggesting possible H-bonding to bases in either bp 4/-4 or 5/-5 (as indicated by the dotted grey lines in Fig. 2b). In 12 of the 20 aligned half-sites (Supplementary Fig. 1b) guanine is found at -5 and in an additional 5/20 half-sites base -5 is thymine, suggesting that H-bond donation from Lys41 to O6 of guanine or O4 of thymine is preferred. His35 and Ser37 are positioned to H-bond to bases in the major groove at positions 6 and -6 respectively. Among MntR recognition sites (Supplementary Fig. 1b), base pair 6/-6 is most commonly an AT base pair (14/20), suggesting preferred H-bonding from His35 to adenine and Ser37 to thymine. However, since both residues are capable of accepting and donating H-bonds, the nature of these interactions is unlikely to be prescriptive in defining a DNA recognition sequence. A hydrophobic interaction between Pro36 and the thymine methyl group at position -7 (19/20 half-sites) does appear to be important in specifying the cognate DNA-binding sites occupied by MntR. The T_-7_ methyl group occupies a pocket lined by the side chains of Thr40 and Val25, in addition to Pro36, suggesting that the C5 methyl group provides uniquely favorable interactions when present at position -7 (Fig. 2). Similar contacts between a proline residue and thymine methyl group, have been identified as playing significant roles in operator recognition in two related proteins, DtxR^17^ and IdeR^18,19^.

Tyr57, which is in the ***β***-turn linking the two strands of the “wing” motif of the winged helix-turn-helix, is positioned to make hydrogen-bonding interactions with base pair edges in the minor groove (Fig. 2 and 3). The side chain lies fully in the minor groove between base pairs 8/-8 and 9/-9 (Fig. 2b). The Tyr57 hydroxyl group could donate a H-bond to either the C2 carbonyl of a pyrimidine or N3 of a purine ring at positions -8 or +9, suggesting that these interactions are not sequence-specific, but rather contribute to the overall affinity of MntR for duplex DNA.

**Figure 3.**
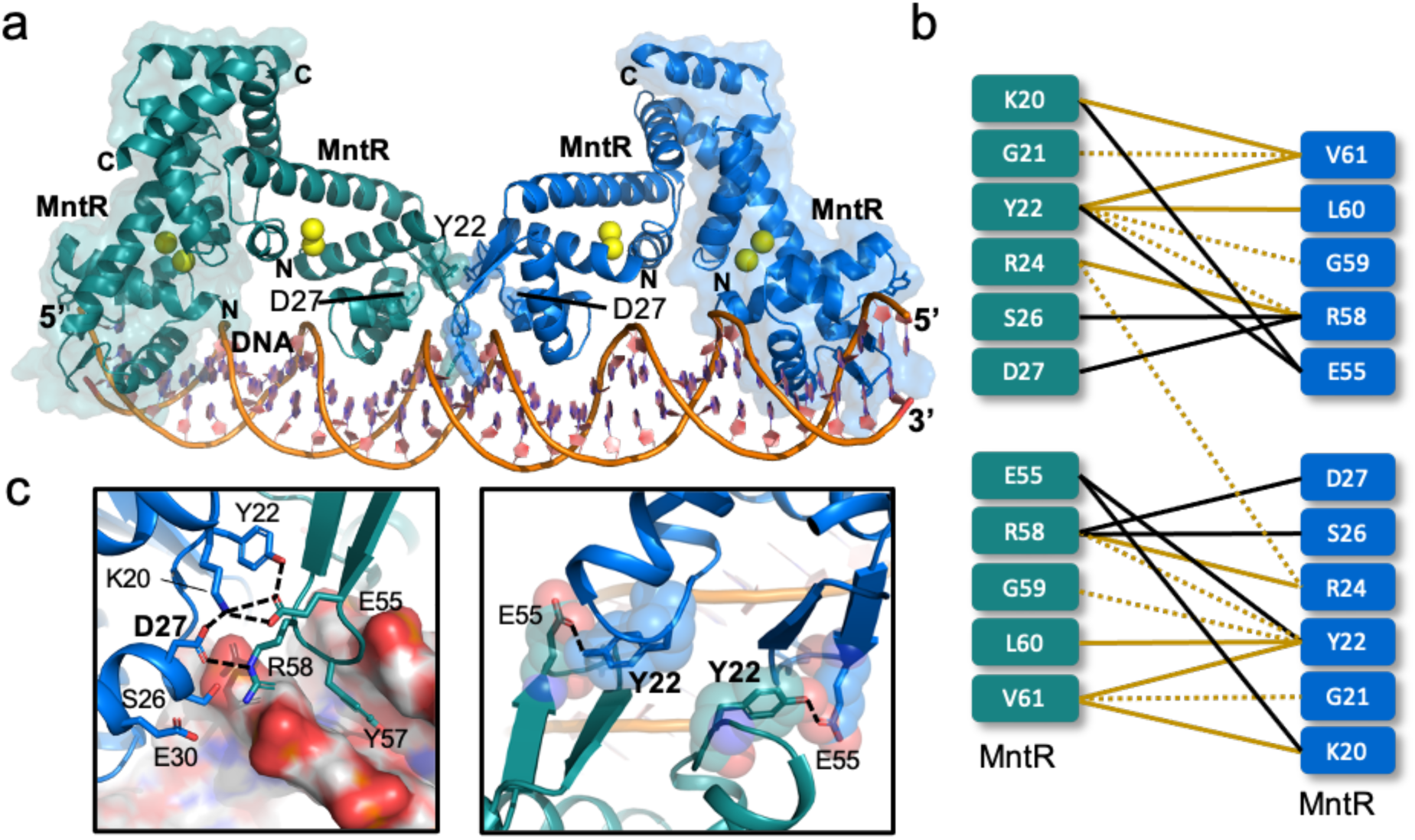
Interdimer interactions observed in the (MntR_2_)_2-_P84 structure. (a) A cartoon representation of the cryo-EM structure of the (MntR_2_)_2_-P84 complex with two MntR dimers highlighted in green and blue, with each dimer bound to 4 Mn^2+^ ions (yellow spheres). The MntR dimers are interacting with a portion of P84 (orange). Tyr22 and Asp27 are highlighted in the cartoon. (b) A schematic representing a summary of interactions observed between the two adjacent dimers of MntR in the (MntR_2_)_2_-P84 structure. Polar interactions such as dipole-dipole interactions, H-bonds and salt-bridges are highlighted by black lines and hydrophobic or vdW interactions are highlighted by orange lines. The dotted lines represent weaker interactions between residues that are separated by a distance ∼4 – 5 Å. Note that the side chain of Tyr22 and Val 61 are interacting with the backbone of Leu60 and Lys20, respectively. (c) Expanded views of the network of interdimer contacts around the conserved Asp27 (left) and Tyr22 (right). The black dotted lines represent polar interactions including salt bridges and H-bonds.

Several other residues contribute to non-specific interactions with phosphates along the MntR-DNA interface (Fig. 2b). Notably, three residues from the wing motif (Tyr54, Lys56, and Arg58) are positioned to contribute H-bonds to phosphate groups at positions -6, -7 and +11 in the DNA duplex. Additionally, the side chains of Arg24, Ser38, Thr40, and Gln44 are near phosphate groups, as are backbone amide nitrogens from Val25, Ser26, His35, and Tyr57 (Fig. 2b). The majority of the protein-DNA interactions observed in the (MntR_2_)_2_-P84 structure involve highly conserved amino acids (Supplementary Fig. 6).

### Protein-protein interactions accompany the formation of the (MntR_2_)_4_-P84 complex

Inspection of the (MntR_2_)_2_-P84 structure reveals a well-defined network of polar and hydrophobic interactions between amino acids at the 974.3 Å^2^ interface between two adjacent dimers of MntR bound to DNA (Fig. 3). Tyr22 is at a dominant position at the interface and participates in interdimer H-bonding with Glu55 while also forming interdimer van der Waals (vdW) interactions with side chain of Val61 and backbone atoms in Gly59 and Leu60. Asp27 is at the heart of a network of polar interactions, forming salt bridges with Arg58 from the second dimer and Lys20 from the same subunit, thus mitigating repulsion between the two residues. Interestingly, Arg24 residues from adjacent MntR subunits stack their guanidinium groups against each other (Supplementary Fig. 7). Simultaneously, each Arg24 forms hydrogen bonds within its own subunit with the carbonyl oxygen of Arg58 and the phosphate group of the nucleotide at position -9. All four arginine residues at the dimer interface form H-bonds with the phosphate groups on the P84. Arginine pairs are relatively common in protein structures, and it has been argued that the polar environment surrounding the arginine residues compensates for what would otherwise be a repulsive interaction^20^. This organization of the positively charged arginine residues at the MntR interdimer interface could be essential to ensure that the arginine side chains are oriented to effectively facilitate polar interaction with either the phosphate backbone of the DNA or with oppositely charged side chains of other residues at the interface.

Several of these residues show strong conservation, for example, Asp27, Arg24, and Arg58 (Supplementary Fig. 6, extended data 1). Sequence alignment of MntR homologs from multiple bacterial species show when tyrosine is present at position 22, glycine is strongly conserved at position 59 (Supplementary Fig. 6, extended data 1). The two interdimer salt bridges observed between largely conserved residues, Asp27-Arg58, and Lys20-Glu55 (Fig. 3b-c), are reminiscent of the two asymmetrical salt bridges identified at the dimer-dimer interface of *E.coli* Zur proteins that bind to the opposite faces of DNA in its regulatory sites^4^.

### MntR interacts cooperatively to bind the *mneP* operator

Given the strong network of interactions between dimers, we were curious to explore the role of these contacts in the association of MntR with DNA in solution. Using mass photometry (MP)^21^ and endogenous tryptophan fluorescence-based size exclusion chromatography (FSEC)^22^ we tested the ability of MntR to form complexes with P84 and a second 84-bp DNA duplex we refer to as C84, which possesses four copies of the consensus MntR binding sequence^8^ in place of sites 1-4 in *mneP* (Supplementary Table 1, Supplementary Fig. 1a). In the presence of 1 mM Mn^2+^, MntR exists as a dimer and is seen as a single peak in the MP experiments at a molecular mass of 44 ± 8 kDa (Fig. 4a), consistent with its predicted mass of 33 kDa. Upon addition of P84, we observed the appearance of a well-defined and reproducible peak at 182 ±18 kDa which corresponds to four MntR dimers bound to a P84 molecule as seen in the cryo-EM generated (MntR_2_)_4_-P84 map. The presence of additional peaks at masses of 111 ± 16 kDa and 156 ± 29 kDa is consistent with two and three MntR dimers on the 84-bp duplex, suggestive of dissociation after dilution to the low complex concentration (16 nM) used in these experiments. In contrast, when MntR is mixed with 1 mM Mn^2+^ and the C84 DNA containing all consensus binding sites the free MntR dimer (44 kDa) disappears and only a single peak at 182 kDa appears, indicative of a stable (MntR_2_)_4_-C84 complex (Fig. 5a). Complementary FSEC experiments conducted on WT MntR indicate the formation of homogeneous complexes of WT MntR with both P84 and C84 as highlighted by the appearance of a symmetric peak at a retention volume smaller than that observed for MntR alone (Fig. 4b and 5b). As for MntR mixed with P84 in FSEC experiments, the lack of multiple species is likely due to the higher concentration of the complex (∼1-8 µM) in this experiment.

**Figure 4.**
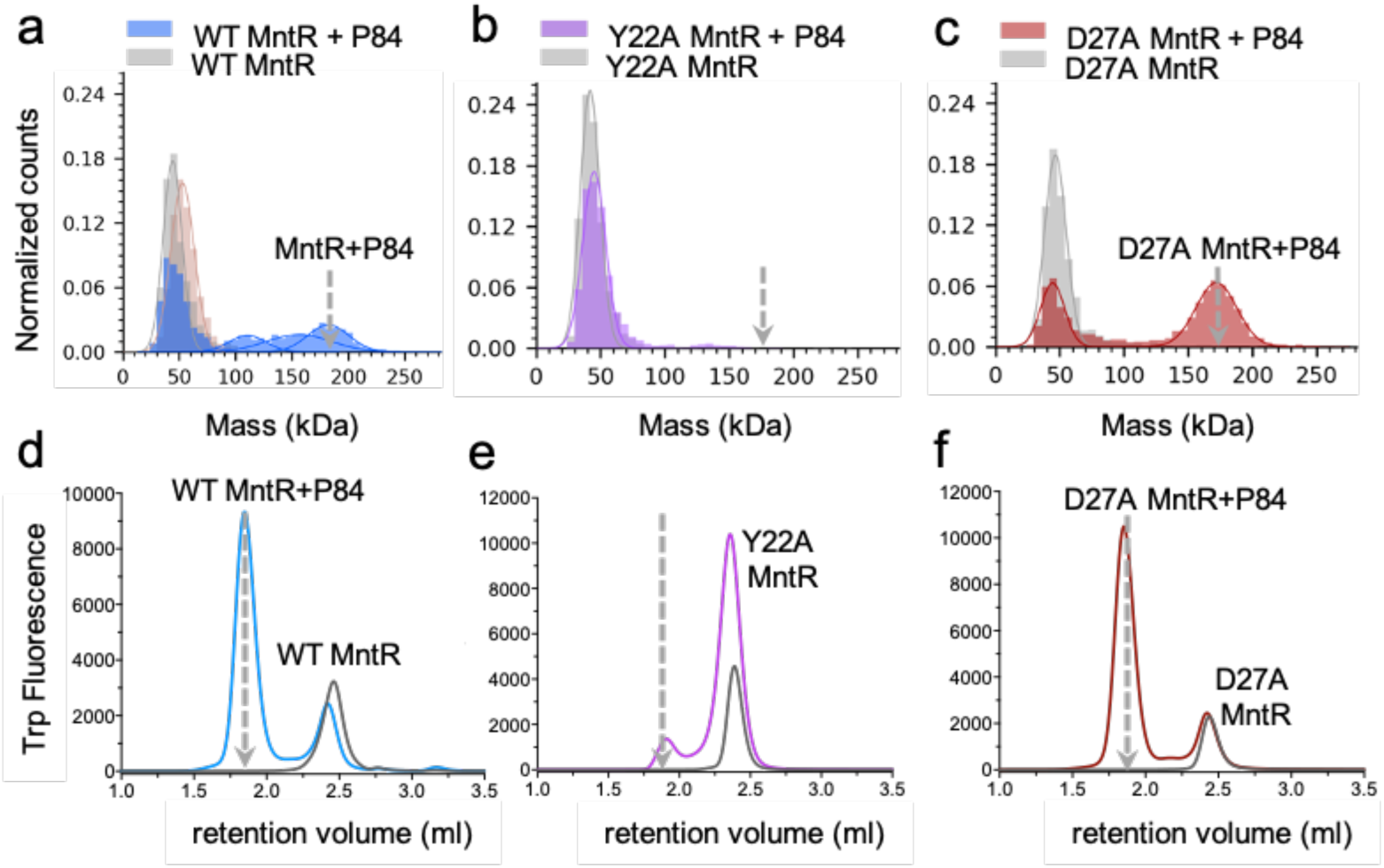
Complex formation of WT and mutant MntR with P84 in solution. Mass photometry data on (a) WT MntR, (b) Y22A MntR and (c) D27A MntR alone (grey) and in complex with the P84 (colored) is presented. The grey broken arrow indicates the expected mass of the (MntR_2_)_4_-P84 complex. (d)-(f) endogenous tryptophan FSEC data was collected on (d) WT MntR, (e) Y22A MntR and (f) D27A MntR alone (grey) and in complex with P84 (colored). The grey broken arrow indicates the expected retention volume of the (MntR_2_)_4_-P84 complex.

**Figure 5.**
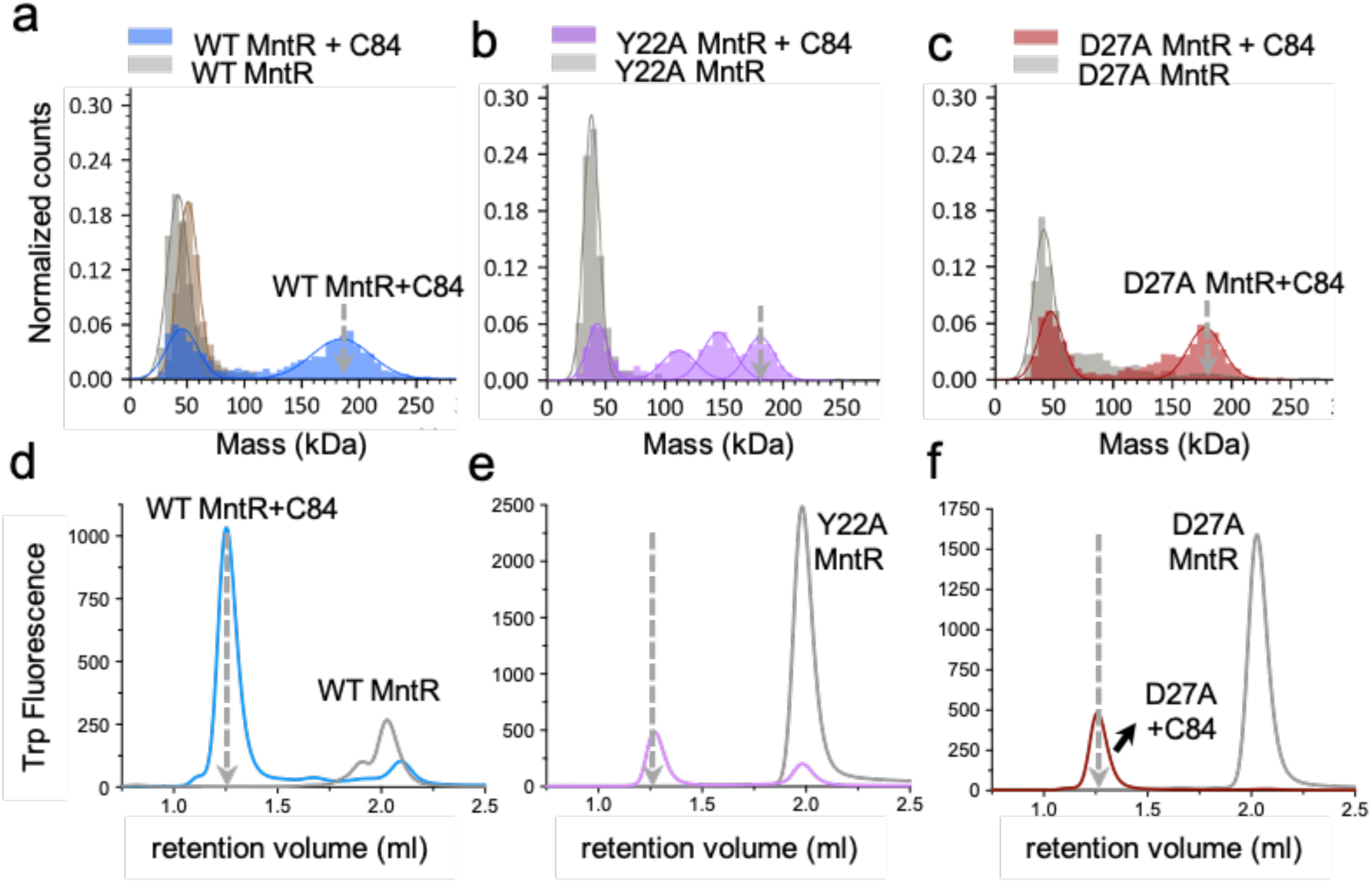
Binding studies on WT and mutant MntR with C84. Mass photometry data on (a) WT MntR, (b) Y22A MntR and (c) D27A MntR alone (grey) and in complex with C84 (colored). The grey broken arrow indicates the expected mass of the (MntR_2_)_4_-C84 complex. (d)-(f) endogenous tryptophan FSEC data on (d) WT MntR, (e) Y22A MntR and (f) D27A MntR alone (grey) and in complex with the C84 (colored). The grey broken arrow indicates the expected retention volume of the (MntR_2_)_4_-C84 complex.

Further FSEC experiments were performed with two 26-bp duplexes. H26 contains the *mntH* operator sequence, while P26 contains the sequence for site 1 from the *mneP* operator (Supplementary Table 1, Supplementary Fig. 1a). FSEC experiments show that WT MntR forms a homogenous complex with H26 at a lower retention volume than for MntR alone but at a larger retention than observed for the MntR-P84 and MntR-C84 complexes, consistent with the smaller size of the 1:1 MntR-H26 complex (Supplementary Fig. 8a). In contrast to these observations, FSEC experiments show that WT MntR does not form a complex with P26, highlighted by the lack of a peak at the lower retention volume (Supplementary Fig. 8a). Site 1 on the *mneP* promoter has a weaker affinity for MntR dimer than the *mntH* operator does. However, WT MntR can bind to P84, a longer sequence that contains all four 18-bp sites (1-4). The necessity of multiple, adjacent MntR binding sites for complex formation suggests that cooperative interactions are required to stabilize the complex of WT MntR with the weaker binding sites used in transcription activation. These results confirm the previous observation that MntR activates expression of MneP with a highly cooperative response to manganese concentration, but not until a higher extracellular concentration of manganese (>10 µM) is reached than that needed to promote repression of MntH expression^8^.

### Mutations to interface residues have significant effects on activation by MntR

Our analysis of residues at the dimer-dimer interface (see above) suggests that Tyr22 and Asp27 are each important to the stability of the functional (MntR_2_)_4_-*mneP* complex. To test that hypothesis, we prepared two MntR variants with Tyr22 and Asp27 substituted with alanine, creating the Y22A and D27A mutants, respectively. We then subjected the variant forms of MntR to the same solution studies as performed with WT MntR using MP and FSEC. In addition, we performed *in vivo* expression tests in *B. subtilis* to compare with our *in vitro* results.

Our *in vitro* studies with the Y22A MntR showed that Tyr22 is indeed an important residue for the cooperative binding of MntR to the *mneP* operator. Molecular photometry experiments indicate that the Y22A MntR does not form a complex with P84, evident by the lack of a peak at 182 kDa (Fig. 4b). Similarly, very little (MntR_2_)_4_-P84 complex is observed in FSEC experiments with Y22A MntR (Fig 4e). In contrast, Y22A MntR successfully forms a complex with C84, behaving similarly to WT MntR in both MP and FSEC experiments (Fig. 4b, e, and Fig. 5b, e). When FSEC is used to probe the formation of protein-DNA complexes between Y22A MntR and either H26 or P26, only the complex with H26 is observed, as was true with WT MntR (Supplementary Fig. 8b). That result is consistent with the formation of the Y22A (MntR_2_)_4_-C84 complex. The mutation of Tyr22 to alanine does not appear to interfere with DNA binding to a single high-affinity site, or four adjacent high-affinity sites, but shows that cooperativity is not available to facilitate the binding of Y22A MntR to the natural DNA sequence in the P84 duplex.

Expression tests in *B. subtilis* reveal that Tyr22 is essential for transcriptional activation but is dispensable for repression. Using ***β***-galactosidase as a reporter gene under the control of the *mntH* and *mneP* promoters, we explored the impact of adding Mn^2+^ to the growth medium on transcription. WT MntR was used as a control, and as expected from a previous study^8^, transcription from *mntH* is repressed in LB medium, with little if any further repression elicited by supplementation with 10 µM Mn^2+^. In contrast, expression from *mneP* is low in LB medium, but is activated when the medium is amended with 100 µM Mn^2+^ (Fig. 6). Supporting the results from MP and FSEC (Supplementary Fig. 8b), Y22A MntR behaves much like WT MntR in repression from *mntH*, which contains a single high-affinity MntR binding site that matches consensus at all highly conserved residues (Supplementary Fig. 1b). However, Y22A MntR does not activate transcription from *mneP*, which contains four weak MntR binding sites. That result coupled with the failure of Y22A MntR to form a complex with P84 suggests that cooperativity is essential in forming the (MntR_2_)_4_-P84 complex. Tyr22 participates in essential interactions in promoting cooperativity between subunits, and its substitution with alanine abolishes those cooperative interactions, without dramatically affecting the ability of the protein to bind high affinity sites in H26 or C84. Mutation of Tyr22 to alanine disrupts essential and conserved interactions at the interface with Glu55, Gly59, and Val61 on the neighboring MntR dimer, thus severely impacting the recruitment of additional MntR dimers to the adjacent sites on the *mneP* promoter and preventing the formation of an active transcription complex.

**Figure 6.**
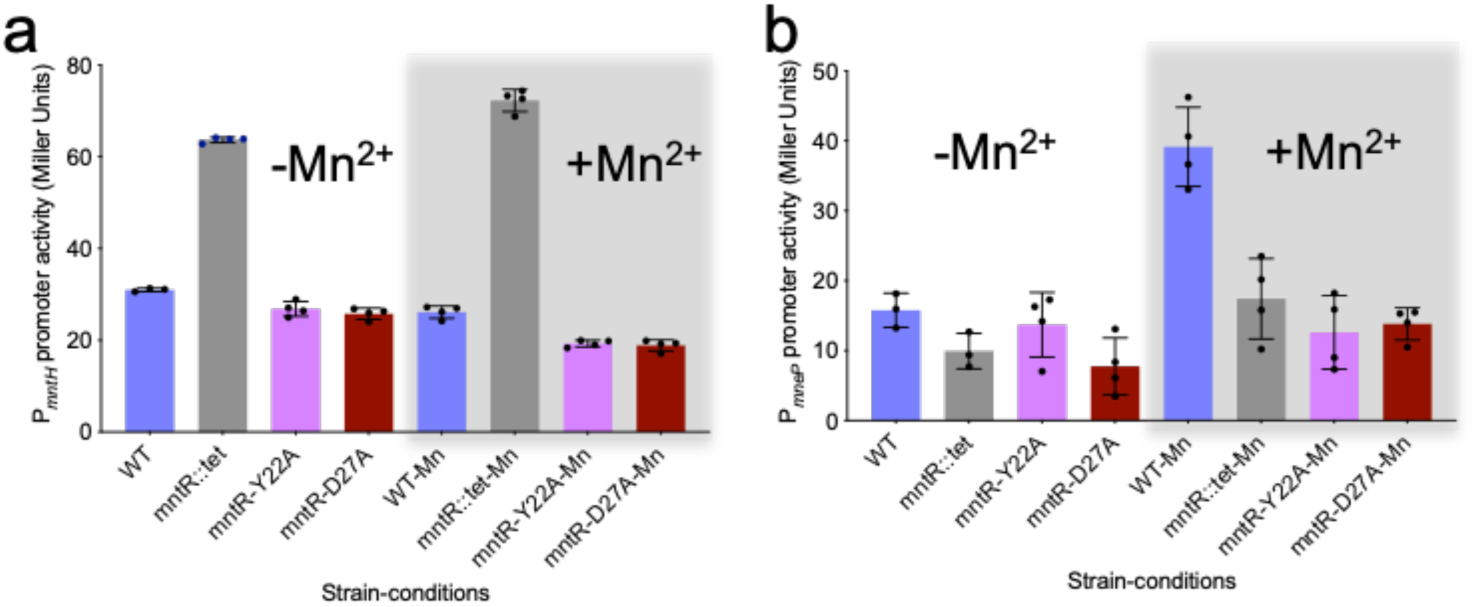
*β*-Galactoside reporter gene assays in *B. subtilis* to study the effect of Y22A and D27A mutations in MntR on the activity of (a) *mntH* and (b) *mneP* promoter. Promoter activities were measured in the presence (grey box) and absence of Mn^2+^ ions. The grey bars in the histograms represent the *mntR::tet* null mutant of *B. subtilis*. Promoter activity was averaged over four replicates as shown and standard deviation is presented as error bars

In contrast with Y22A MntR, the D27A MntR mutant forms (MntR_2_)_4_-P84 (Fig. 4c, f) and (MntR_2_)_4_-C84 (Fig. 5c, f) complexes. It also binds the H26 duplex but fails to form a complex with P26 in the FSEC experiment (Supplementary Fig. 8c). The presence of a single peak at ∼182 kDa in MP data (Fig. 4c), indicates that D27A MntR forms a higher affinity complex with P84 than WT MntR (Fig. 4a). Despite being involved in polar interactions with a network of amino acids across the MntR dimer-dimer interface, Asp27 is not essential for MntR binding to the *mneP* operator sequence. The *in vivo* reporter gene assay, however, shows that D27A MntR does not activate transcription from the *mneP* promoter, despite being competent at repression from the *mntH* promoter (Fig. 6). Hence, Asp27 does not seem to play an essential role in DNA-binding by MntR but is important for activation of transcription. The fact that D27A MntR binds P84 better than WT MntR (as observed in MP experiments) may relate to the loss of a negatively charged residue at the protein-DNA interface. The substitution of Asp27 by alanine appears to increase the affinity to DNA, which might compensate for any decrease in cooperativity. The failure of D27A MntR to activate transcription from the *mneP* promoter in 100 µM Mn^2+^ may reflect a loss of stabilizing interactions between MntR and RNA polymerase.

The network of interactions involving Asp27 at the MntR dimer-dimer interface in our (MntR_2_)_2_-P84 structure provides a plausible explanation for these results (Fig. 3). Asp27 forms a salt bridge with the side chain of Lys20 on the same MntR and Arg58 on the neighboring MntR dimer (Fig. 3c). Arg58, in turn, interacts with the phosphate group of nucleotide +11 in the MntR-recognized half site (Fig. 2b). Mutation of Asp27 to alanine allows the conserved Arg58 to form stronger interactions with the backbone phosphate group on the DNA duplex, potentially increasing MntR’s affinity for P84, consistent with mass photometry experiments (Fig. 4). Additionally, mutation of Asp27 to a small, non-polar residue like alanine also allows Lys20 to interact with and further stabilize the network of interactions with the Tyr22 side chain on the same MntR and the carboxylate of Glu55 on the adjacent MntR dimer, consistent with in-vitro binding studies to the P84. Additionally, the RNAP binding site overlaps the MntR-dimer binding site 1 on the *mneP* promoter sequence (Fig. 1a). Hence, Asp27 on the external MntR subunit at site 1, could be involved in binding to RNAP and recruiting it to the *mneP* promoter. Hence, Asp27 could be necessary for activation of transcription *in vivo* but not critical for binding of MntR to the *mneP* promoter.

### Conclusion

MntR from *B. subtilis* displays distinctive differences in its function as a repressor and activator. MntR forms 1:1 complexes with the high-affinity *mntH* and *mntA* operators^23^ to repress expression of uptake transporters without the need for interdimer contacts. However, to activate expression from the *mneP* promoter, the binding of four MntR dimers, as seen in our cryo-EM structure, depends on cooperative protein-protein interactions, as is evident from solution experiments and *in vivo* expression tests described above.

Similar to *B. subtilis*, the regulation of manganese uptake and efflux in *E. coli* is controlled by the dual activity of an MntR homolog. At high cellular Mn availability, the MntR from *E. coli* represses the expression of an uptake transporter, MntH, and a small peptide implicated in Mn homeostasis, MntS^24–27^. Also, at high Mn availability, MntR appears to activate the expression of an efflux transporter, MntP^25^. Interestingly, the *mntS* and *mntP* genes are under the control of paired MntR binding sites. In the case of *mntS,* two binding sites overlap with the promoter. Assuming an 18-bp recognition sequence of two 9-bp inverted repeats, the spacing of the MntR binding sites in the *mntS* operator are identical to those observed in the *mneP* and *mneS* operators, suggesting a common means of achieving cooperative binding for *E. coli* and *B. subtilis* MntR homologs. On the other hand, the two MntR binding sites upstream of the *mntP* gene are well separated from each other (over 20 bp) and distant from the promoter sequence (∼170 bp), suggesting different mechanisms of activation by *E. coli* MntR and *B. subtilis* MntR^25^.

Our cryo-EM structures, along with solution studies and *in vivo* transcription assays, show that high cooperativity in the binding of MntR dimers to the *mneP* promoter driven by interdimer contacts is necessary to activate transcription of *mneP* in steep response to increase in concentration of Mn^2+^ ions above 10 *μ*M. Cooperativity is a well-known feature in protein-DNA interactions and has been extensively studied, starting with the lambda repressor^28^.

Metalloregulation, in several instances, also relies on cooperativity between multiple repressor proteins binding to a regulatory region on DNA. Zur^4^ shows a high degree of cooperativity between two dimers that bind on opposite faces of the operator sequence as does the iron-responsive regulator DtxR^17^. The Mn-responsive regulatory protein from *S. mutans*, SloR, binds to three adjacent sites upstream of an operon encoding an ABC transporter with positive cooperativity to repress transcription^29^. Cooperativity increases the steepness of the response curve to increasing metal ion availability in the cell and may be particularly desirable when increasing cellular metal ion availability beyond a certain concentration has significant negative consequences. Our study suggests that high cooperativity in MntR binding to the efflux promoter is necessary for the activation of transcription. How this cooperative binding recruits RNAP is still unclear and further structural work is needed to understand the full mechanism of transcription activation by MntR.

## METHODS

### Expression of MntR and its mutants

WT, Y22A, and D27A MntR proteins was expressed from the pSMT3 vector, a derivative of pET28b Kan^r^, which encodes a His_6_ tagged SUMO (small ubiquitin-like modifier) domain downstream of the T7 promoter and the *lac* operator^30^. The WT and mutant plasmids were constructed via site-directed PCR mutagenesis (primers are listed in Supplementary Table 1). All plasmids were verified by sequencing and then transformed into BL21(DE3)NiCo cells from New England Biolabs for expression. Expression of WT and mutant MntR was induced by adding 0.5 mM IPTG to cultures grown in Luria Broth at 37°C with shaking to an OD600 of 0.6. After three more hours of incubation, cells were harvested via centrifugation and the cell paste was stored at -80°C.

### Purification of MntR

Cell paste was resuspended in chilled Buffer A (25 mM HEPES, pH 7.5, 300 mM NaCl, 10 mM imidazole, 5% glycerol) with the addition of 6 mg of DNAase, 1 mM MgCl_2_, 100 µg/ml lysozyme, and cOmplete EDTA-free protease inhibitor cocktail tablet. After cell lysis by sonication and clarification by centrifugation, the supernatant was loaded onto a 5 ml column of TALON® Metal Affinity Resin (TAKARA) equilibrated with Buffer A. The column was washed with 10 CV of 10 mM imidazole buffer, followed by 10 CV of 20 mM imidazole buffer, and then eluted with 5 CV of 300 mM imidazole buffer. Fractions containing mutant SUMO-MntR were pooled, incubated overnight with ULP-1 protease (Ub1-specific protease-1 from *Saccharomyces cerevisiae*) at a protein-to-protease ratio of 1:20, and dialyzed against Buffer A with 1mM β-mercaptoethanol to cleave the His_6_ tagged SUMO domain from MntR protein.

Cleaved MntR was loaded onto a 5 ml column of Ni-NTA His•Bind® Resin (Millipore Sigma) equilibrated with Buffer A and then eluted with 40 mM imidazole. The column was then washed with 300 mM imidazole to remove the remaining cleaved His_6_-SUMO tag and ULP-1 protease. MntR-containing fractions were pooled, concentrated, and dialyzed against storage buffer (25 mM HEPES pH 7.4, 300 mM NaCl, 10% (v/v) glycerol). The next day the dialyzed protein was aliquoted and flash-frozen for storage at -80°C.

### Preparation of DNA duplexes

All synthetic oligonucleotides used in this study were purchased from either IDT or Oligos Etc. Inc. and were dissolved in an annealing buffer (25 mM HEPES pH 7.4 and 50mM NaCl) to prepare 40 µM-1000 µM solutions of single-stranded DNA. Solutions of complementary strands were mixed in equal molar concentrations (20 µM-500 µM) and annealed via heating to 90 °C and slowly cooled down to room temperature at the rate of 1 °C per 3 minutes. The resulting DNA duplexes were verified by running on a polyacrylamide gel (data not shown) and stored at - 20°C.

### Cryo-EM sample preparation, data collection, and processing

Samples were applied to glow-discharged grids and plunge-frozen in liquid ethane. Then, the grids were clipped and sent to the Pacific Northwest Center for Cryo-EM (PNCC) for imaging. The samples consisted of the following: 8 μM MntR, 1 μM P84 DNA duplex, 20 mM HEPES pH 8.0, 0.05% Tween20, 1 mM MnCl_2_, and 500 mM NaCl. The grids used were Quantifoil R2/1 copper holey carbon grids with 200 mesh. The glow discharge parameters were as follows: 0.38 mBar, 15 mA negative set, 1 minute glow, 30-second hold. Sample grids were screened using a Talos Artica at PNCC. Movies were collected on a Titan Krios III at 300 kV in multishot mode, where 3 images were collected per hole and 9 holes per stage move. Pixel size was 0.826 Å/pixel and super-resolution pixel size was 0.426 Å/pixel. The total dose was 50 e-over 50 frames. The defocus range was -0.8 to -2.2. All processing steps were conducted using cryoSPARC^31^. After data collection, 7215 super-resolution micrographs were pre-processed via motion correction and local CTF correction, followed by micrograph curation, resulting in 6532 accepted micrographs. An initial volume representing two MntR dimers bound to DNA was generated from blob picking and iterative 2D refinement. Template picking and subsequent 2D and 3D iterative refinement yielded a final particle stack of 194,072 particles (Supplementary Fig. 2). The resolution of the final map of two MntR dimers bound to P84 ((MntR_2_)_2_-P84 complex) was estimated to be 3.09 Å, as measured by gold-standard Fourier shell correlation (GSFSC) at a threshold of 0.143 (Supplementary Fig. 3). To obtain the map of four MntR dimers bound to P84 ((MntR_2_)_4_-P84 complex), the (MntR_2_)_2_-P84 complex map was used to generate template picks with an enlarged box size, followed by iterative 2D and 3D refinement, resulting in 228,659 particles. The final refinement yielded a (MntR_2_)_4_-P84 map with a nominal resolution of 4.17 Å.

### Model Building

Models were fit to both the (MntR_2_)_2_-P84 and (MntR_2_)_4_-P84 maps using the known crystal structures of MntR (PDB: 2F5C)^11^ and of the MntR homolog, IdeR, bound to DNA (PDB: 1U8R)^19^. The chosen crystal structure of MntR, among the many deposited of MntR-metal complexes, has the advantage of including the flexible wing region in the winged helix-turn-helix motif (residues 52-62) and the full C-terminal alpha helix, spanning residues 124-142. A starting model to fit each MntR dimer bound to DNA was created by aligning the MntR dimer to the IdeR dimer in the presence of the DNA duplex. The lower resolution (MntR_2_)_4_-P84 map provides density for most of each MntR chain, from residues 3-142, while (MntR_2_)_2_-P84 map lacks density for the C-terminal 6-7 residues in all four MntR subunits (Supplementary Fig. 3). The “wing” (residues 52-62) of the winged helix-turn-helix is often disordered in crystal structures of MntR without DNA^11^ but is clearly visible interacting with DNA in both maps. Real space refinement of the model to the maps was performed in Phenix^32^ with the inclusion of secondary structure restraints derived from crystal structures of MntR-Mn^2+^ complexes (PDB: 2F5D and 2F5C), base-pairing and stacking restraints and geometric restraints related MntR-metal interactions, via residues Asp8, Glu11, His77, Glu99, Glu102 and His103. Rebuilding was performed in Coot^33^ and Isolde^34^. The (MntR_2_)_2_-P84 model includes independent xyz and ADP refinement of all four MntR subunits, while NCS restraints were used for the eight MntR subunits in the final xyz refinement of the (MntR_2_)_4_-P84 complex, which was followed by a final round of independent ADP refinement. The stereochemical quality of the models was evaluated using Molprobity^35^ as implemented in Phenix (Supplementary Table 2).

### Mass Photometry

All mass photometry measurements were performed on a TwoMP0224 mass photometer (Refeyn Ltd). MntR-DNA complex samples were prepared at a concentration of 1.0 µM of the appropriate DNA duplex (P84 or C84) and an excess of ∼8.0 µM MntR in Buffer B (25 mM HEPES pH 7.4, 300 mM NaCl, and 1 mM MnCl_2_). The high micromolar concentrations of MntR and DNA duplexes were chosen to be about x1000 fold above the apparent dissociation constant (*K_d_*) ∼ 1.4 nM^8^ of WT MntR for the *mneP* promoter DNA in the presence of Mn^2+^ ions. The complex samples were prepared and stored on ice for at least an hour before the MP experiments and the samples were further diluted 10-fold in buffer B immediately before the MP measurements were performed.

Precut six-well silicone gaskets were positioned atop precleaned and poly-L-lysine coated glass coverslips to accommodate six samples per coverslip. The poly-L-lysine coating helps DNA adhere to the glass coverslips. These coverslips were subsequently positioned on the stage of a Two-MP mass photometer instrument. Utilizing the lateral control button within the software, the first well was maneuvered over the objective, and 18 µl of PBS buffer was dispensed into one of the gaskets for MP measurement. Subsequently, 2 µl of the diluted MntR-DNA sample was added and mixed well to achieve a final concentration of ∼16 nM MntR. Sample binding to the poly-l-lysine coverslip was monitored via a one-minute-long movie that was recorded using the acquisition software AcquireMP (AMP) version 2024 R1.1. Standard proteins, such as *β*-Amylase (BAM; mass: 56 kDa, 112 kDa, 224 kDa) and Thyroglobulin (TG; mass: 670 kDa), underwent measurement in a similar manner as the samples to establish a calibration curve on the same day. Recorded movies were analyzed using the Discover MP (DMP) 2024 R1.0 software, ensuring a mass error of less than 5%. A linear calibration curve was constructed using BAM and TG movies in the DMP software, associating the proteins’ masses with the Ratiometric contrast values and subsequently applied to the sample proteins to ascertain their molecular mass in kDa.

### Fluorescence-based size exclusion chromatography (FSEC) measurements

MntR (WT and mutants) and DNA (C84, P84, P26, and H26) complex samples were prepared at a micromolar concentration as described in the mass photometry section above and were analyzed by size exclusion chromatography coupled with fluorescence detection (FSEC)^22^. Samples were transferred into the wells of a 96-well sample block maintained at 4 ℃. A Waters ACQUITY Arc Bio UPLC/SHPLC system with a fluorescence detector was used to apply 30 μL - 50 μL of each sample to a Superose6 or Superdex 200 Increase GL 5/150 column equilibrated in Buffer B. Each sample was analyzed for tryptophan fluorescence (excitation at 280 nm and emission at 350 nm). Each chromatographic run was over two-column volumes (∼20 minutes) at a flow rate of 0.3 ml/min. The data was analyzed and plotted using Prism (GraphPad).

### B. subtilis bacterial strain construction and growth conditions

The bacterial strains of *B. subtilis* used in this study are listed in Supplementary Table 3 and primers are listed in Supplementary Table 1. The *mntR* point mutations were created using CRISPR-based mutagenesis. To this end, we used long-flanking homology (LFH) PCR to generate allelic variants of *mntR* with sufficient upstream and downstream homology. These repair template PCR fragments were cloned into pAJS23 plasmid, which expresses an*erm*-directed guide RNA^36^. The recombinant plasmids were transformed into *mntR*::*erm B. subtilis* strains at 30°C and transformants were selected onto LB (Lennox L Broth, RPI) containing kanamycin (15mg/mL) and 0.2% mannose. Plasmid was cured at 45°C by repeated passaging of transformants on LB media. The erythromycin and kanamycin negative strains were selected for Sanger sequencing for further confirmation. The components of minimal media were the same as previously described^36^. Whenever applicable, we used MLS (erythromycin (1μg/mL) plus lincomycin (25μg/mL)).

### Promoter β-galactosidase assay in B. subtilis

A small amount of respective bacterial strains was taken from a plate and inoculated into 5 mL of LB broth. Cultures were grown to an OD600 nm of 0.4 to 0.6 at 37°C with shaking. The strains containing P*_mntH_*-lacZ and P*_mneP_*-lacZ reporter fusions in WT, *mntR*::*tet*, and *mntR* (Y22A or D27A) were grown in Luria Bertani broth (LB) to a mid-log phase (OD600nm =0.4). Cells of P*_mntH_*-lacZ fusions were treated with and without 10 µM Mn^2+^ and P*_mneP_*-lacZ fusions were treated with and without 100 µM Mn^2+^ ions. Cells were incubated aerobically for 15 min, harvested by centrifugation for 5 min at 8000 rpm, and resuspended in Z buffer containing freshly added 400 nM DTT. 150 µl of cell suspension was used to measure OD 600nm and was incubated with 30 µl of lysozyme solution (10 mg/ml). Lysis was conducted at 37 °C for 15 min and immediately 20 µl of o-nitrophenyl-***β***-d-galactopyranoside (ONPG) solution was added to a concentration of 4 mg/ml. The changes in absorbance at 420 and 550 nm were monitored for 60 min with 3 min intervals using BioTek plate reader. Promoter activities were calculated by measuring ONP production using the formula 1000*[OD420-(1.75*OD550)]. The slope was calculated and normalized to OD600 (slope/OD600), where this quotient provides the promoter activity in Miller units (MU). Replicate data for promoter activity was averaged and standard deviation was taken. Error bars corresponding to standard deviation were added.

### Sequence alignment

Sequence analysis of MntR and its homologs was based on the results of a BlastP^38^ search against the *B. subtilis* MntR sequence using the refseq_select database^39^, extending the results to an E value of 1e-15. That set of sequences was further edited to remove partial sequences. Following that edit, 2334 sequences remained. To analyze dimer-dimer interactions, a spreadsheet was created (supplemental data 1) that selected as individual columns the residues at positions 20, 22, 26, 27, 30, 55, and 58. The Weblogo for the alignment was created using the Berkeley Weblogo server^40^ (https://weblogo.berkeley.edu/logo.cgi).

## Abbreviations

WT: Wild Type
MntR: Manganese transport regulator
Cryo-EM: cryogenic electron microscopy
MP: mass photometry
FSEC: Fluorescence-based size exclusion chromatography
GSFSC: Gold-standard Fourier shell correlation

## Acknowledgments

We thank R. M. Haynes, C. Yoshioka, and C. López at PNCC for access and microscopy assistance. We thank Dr. Isabelle Baconguis and Dr. Steve Mansoor at Oregon Health & Science university for the use of their equipment. We thank Adam Oken at Oregon Health & Science university for helpful discussion of processing cryo-EM datasets. We thank Refeyn Inc. for the use of their mass photometer. We thank Saroj Mahato for assistance with the CRISPR mutagenesis. This work was supported by National Institutes of Health grant R35GM122461 (JDH). The content is solely the responsibility of the authors and does not necessarily represent the official views of the National Institutes of Health. A portion of this research was supported by NIH grant U24GM129547 and performed at the PNCC at OHSU and accessed through EMSL (grid.436923.9), a DOE Office of Science User Facility sponsored by the Office of Biological and Environmental Research.

## Statements and Declarations

### Author Contributions

All authors contributed to the study conception and design, material preparation, data collection and analysis. SA supervised the studies. The first draft of the manuscript was written by AG, HS and SA, and all authors commented on previous versions of the manuscript. All authors read and approved the final manuscript.

### Data Availability

The model and EM maps for (MntR_2_)_4_-P84 (PDB ID 9C4D, EMD-45182) and (MntR_2_)_2_-P84 (PDB ID 9C4C, EMD-45181) complex have been deposited with the Protein Data Bank. Other datasets generated and/or analyzed during the study are not publicly available because they have not been uploaded to any public repository, but are available from the corresponding author on reasonable request.

### Conflict of Interest

The authors declare no relevant financial or non-financial competing interests.

## Notes

### Competing Interest Statement

The authors have declared no competing interest.

### Summary of Updates

The acknowledgment section has been updated.

