## Supplementary Tables and Figures for "Structural basis for transcription activation through cooperative recruitment of MntR"

**Supplementary Table 1: Oligonucleotides used in this study**

| DNA Duplexes created | Primer Name | Sequence |
|---|---|---|
| **P84** | MneP84F | cctttttagcatagctccaactttttttctgtcaccttattta ttagtaaacaggaaacaacgttgctatagacccactcgaag |
| | MneP84R | cttcgagtgggtctatagcaacgttgtttcctgtttactaata aataaggtgacagaaaaaaagttggagctatgctaaaaagg |
| **C84** | Cons84F | ccttttttgccttaaggaaacttttttgccttaaggaaactta tttgccttaaggaaactactttgccttaaggaaactcgaag |
| | Cons84R | cttcgagtttccttaaggcaaagtagtttccttaaggcaaata agtttccttaaggcaaaaaagtttccttaaggcaaaaaagg |
| **H26** | MntH26F | GATAATTTGCCTTAAGGAAACTCTTC |
| | MntH26R | CTATTAAACGGAATTCCTTTGAGAAG |
| **P26** | P26-FOR | CAACGTTGCTATAGACCCACTCGAAA |
| | P26-REV | TTTCGAGTGGGTCTATAGCAACGTTG |
| | MntR_Y22A_FOR | AGAAAAAGGAgcgGCACGAGTTTCC |
| | MntR_Y22A_REV | TCAATCAGCATATAAATCTGTTC |
| | MntR_D27A_FOR | ACGAGTTTCCgcgATCGCGGAAG |
| | MntR_D27A_REV | GCATATCCTTTTTCTTCAATCAG |
| | SfiI-BSU24520_UF1 | AAGGCCAACGAGGCCCTCGGTCTGGACGCTTACTTTGCCATCC CG |
| | SfiI-SBU24520_DR1 | AAGGCCTTATTGGCCCGGCAATAATGGCGCCGGCGCTTG |
| | BSU24520-Y22A_DF1 | GCTGATTGAAGAAAAAGGAGCGGCACGAGTTTCCG |
| | BSU24520-Y22A_UR1 | CGGAAACTCGTGCCGCTCCTTTTTCTTCAATCAGC |
| | BSU24520-D27A_DF1 | GCACGAGTTTCCGCGATCGCGGAAGCATTG |
| | BSU24520-D27A_UR1 | CAATGCTTCCGCGATCGCGGAAACTCGTGC |

### Supplementary Table 2. Cryo-EM collection, refinement, and validation statistics

| | **$(MntR_2)_2$-P84 DNA** (EMD-45181) (PDB: 9C4C) | **$(MntR_2)_4$-P84 DNA** (EMD-45182) (PDB: 9C4D) |
|---|---|---|
| **Data collection and processing** | | |
| Magnification (kx) | 105 | 105 |
| Voltage (kV) | 300 | 300 |
| Electron exposure (e–/Å$^2$) | 50 | 50 |
| Movie frames | 50 | 50 |
| Defocus range (µm) | -0.8 to -2.2 | -0.8 to -2.2 |
| Pixel size (Å) | 0.426 super-resolution | 0.426 super-resolution |
| Symmetry imposed | C1 | C1 |
| Initial micrographs (no.) | 7,213 | 7,213 |
| Final micrographs used (no.) | 6,532 | 6,532 |
| Initial particle images (no.) | 5.9 million | 778,718 |
| Final particle images (no.) | 194,072 | 228.659 |
| Map resolution (Å) | 3.09 | 4.17 |
| FSC threshold | (0.143) | (0.143) |
| **Refinement** | | |
| Map sharpening *B* factor (Å$^2$) | 91.6 | 143.4 |
| **Model composition** | | |
| Non-hydrogen atoms | 6076 | 12493 |
| Protein Residues | 542 | 1120 |
| Nucleotide | 77 | 154 |
| Ligands | 8 | 16 |
| Waters | 0 | 0 |
| *B* factors (Å$^2$) | | |
| Protein | 0/77.91/27.76 | 22.87/300.60/161.10 |
| Metal Ions | 47.87/92.29/59.53 | 127.36/291.29/191.93 |
| Nucleotide | 0/127.81/53.98 | 108.01/437.13/201.13 |
| R.m.s. deviations | | |
| Bond lengths (Å) | 0.004 (0) | 0.002 (0) |
| Bond angles (°) | 0.711 (7) | 0.480 (1) |
| Validation | | |
| MolProbity score | 0.98 | 1.66 |
| Clash score | 2.07 | 6.93 |
| Poor rotamers (%) | 0 | 0 |
| Ramachandran plot | | |
| Favored (%) | 99.25 | 99.28 |
| Allowed (%) | 0.75 | 0.72 |
| Disallowed (%) | 0 | 0 |

**Supplementary Table 3.** ***B. subtilis*** **strains used for the study**

| Strain or plasmid | Genotype | Construction | Reference or source |
|---|---|---|---|
| HB17797 | *thrC::P*$_{mneP}$*-lacZ* MLS, (CU1065) | Lab stock | [8]PMID: 27748968 |
| HB19513 | *amyE*::P$_{mntH}$ SPβ 7510 P$_{mntH}$*-cat-lacZ* MLS | Lab stock | [8]PMID: 27748968 |
| HBRR16 | *mntR*-Y22A thrC::P$_{mneP}$*-lacZ* MLS, (CU1065) | LFH, mntR-Y22A+pAJ23→ *mntR*::*erm* (CRISPR); *thrC*::P$_{mneP}$*-lacZ gDNA transformation* | This study |
| HBRR25 | *mntR*-D27A *thrC*::P$_{mneP}$*-lacZ* MLS, (CU1065) | LFH, mntR-D27A+pAJ23→ *mntR*::*erm* (CRISPR); *thrC*::P$_{mneP}$*-lacZ gDNA transformation* | This study |
| HBRR22 | *mntR*-Y22A *amyE*::P$_{mntH}$*-cat-lacZ* MLS, (CU1065) | mntR-Y22A+pAJS23→ *mntR*::*erm* (CRISPR); *amyE*::P$_{mntH}$*-cat-lacZ* gDNA transformation | This study |
| HBRR07 | *mntR*-D27A amyE::P$_{mntH}$*-cat-lacZ* MLS, (CU1065) | mntR-D27A+pAJS23→*mntR*::*erm* (CRISPR); ; *amyE*::P$_{mntH}$*-cat-lacZ* gDNA transformation | This study |

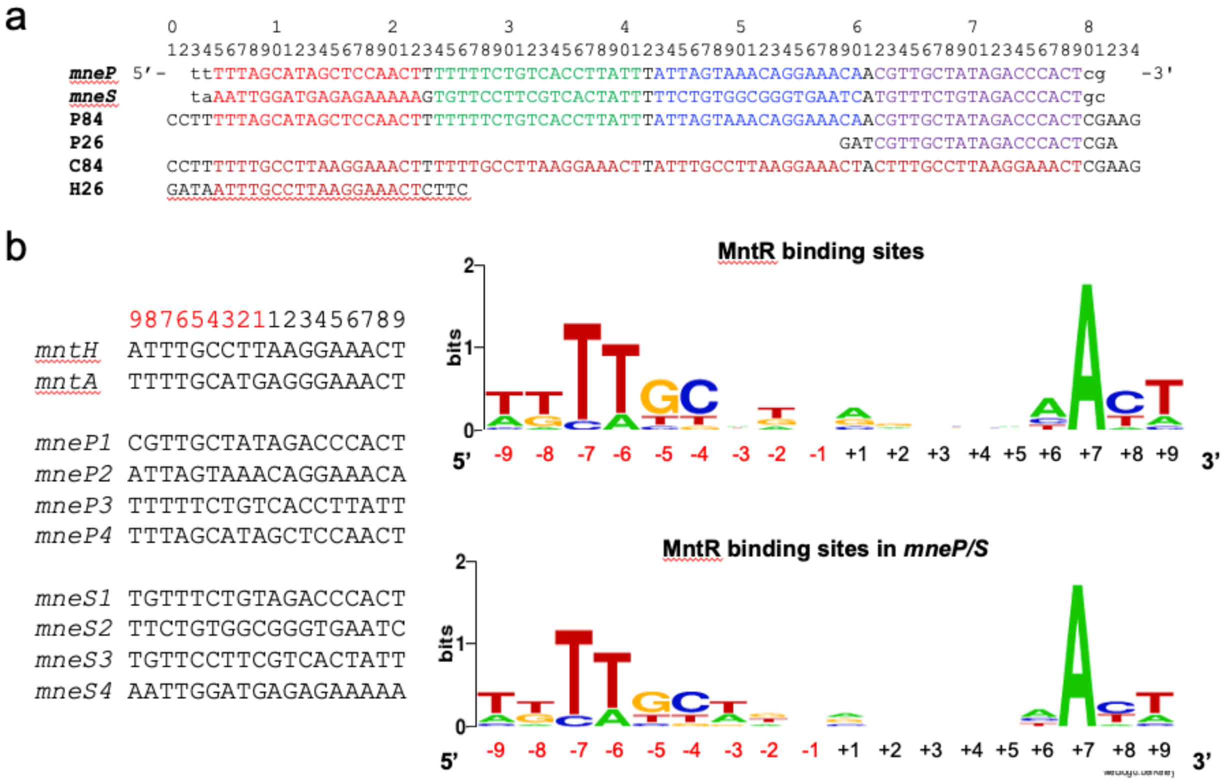


**Supplementary Figure 1. MntR binding site sequence alignments.** (a) *mneP* and *mneS* promoter sequences containing all the four observed binding sites for MntR dimers (site 1 (purple), site2 (blue), site 3 (green), and site 4 (red)). The panel also shows P84 containing the *mneP* promoter sequence, C84 containing all four consensus sites (orange), P26 containing the mneP promoter site 1 (purple), and H26 containing the mntH promoter site (bold). (b) MntR binding sites from *mntA, mntH, mneP* and *mneS* promoter sequences were aligned (left) and a sequence logo was created (right) using Weblogo (http://weblogo.berkeley.edu/logo.cgi). The overall height of each stack indicates the sequence conservation at that position (measured in bits), whereas the height of each letter within the stack reflects the relative frequency of the corresponding base at that position.

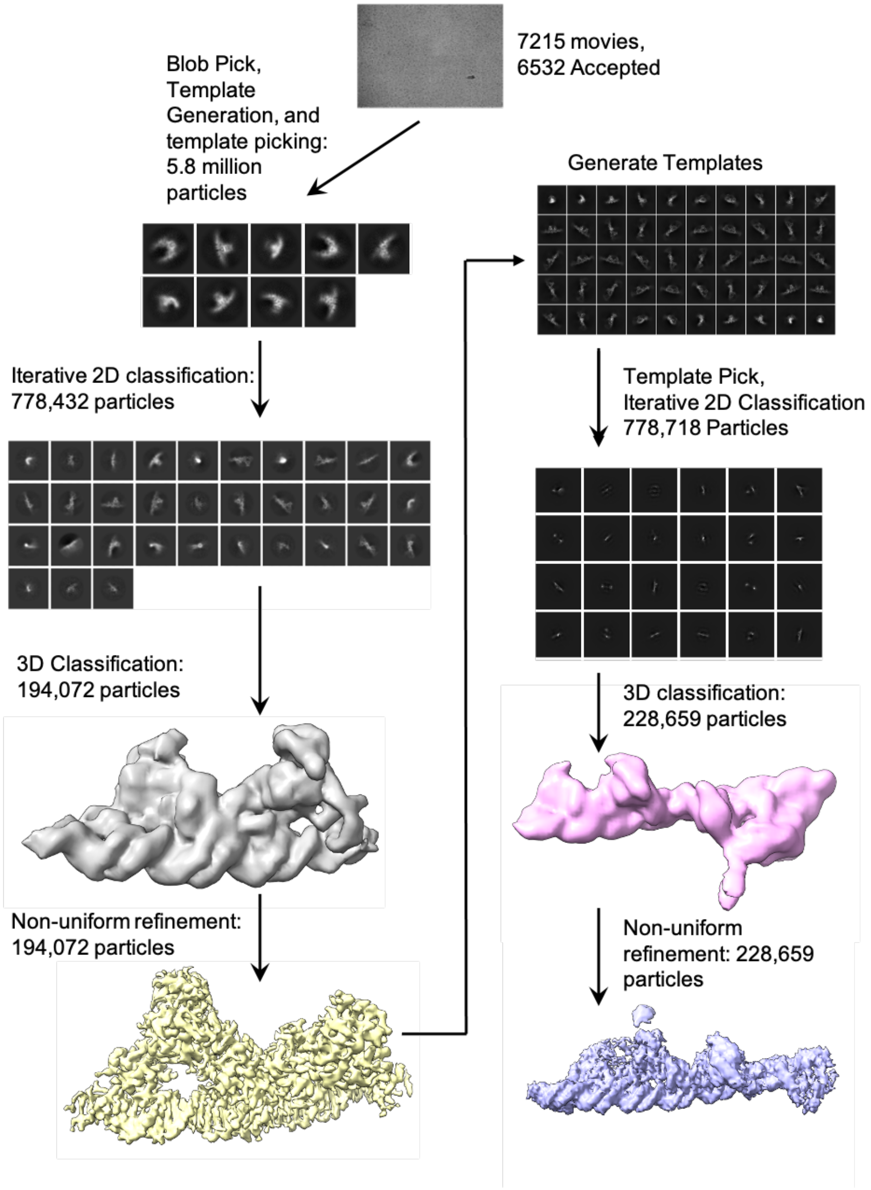


**Supplementary Figure 2. Cryo-EM processing pipeline.** 7215 movies were preprocessed via local motion and contrast-transfer function correction, followed by micrograph curation, resulting in 6532 movies. After blob picking to generate templates for particle picking, particles were obtained and iteratively refined via 2D and 3D classification, resulting in reconstruction of $(MntR_2)_2$-P84. To obtain the complete $(MntR_2)_4$-P84 DNA, the obtained map was used to generate templates for template picking. The resulting particles were extracted at a larger box size to include the entirety of the complex. The final map is produced through iterative 2D and 3D classification.

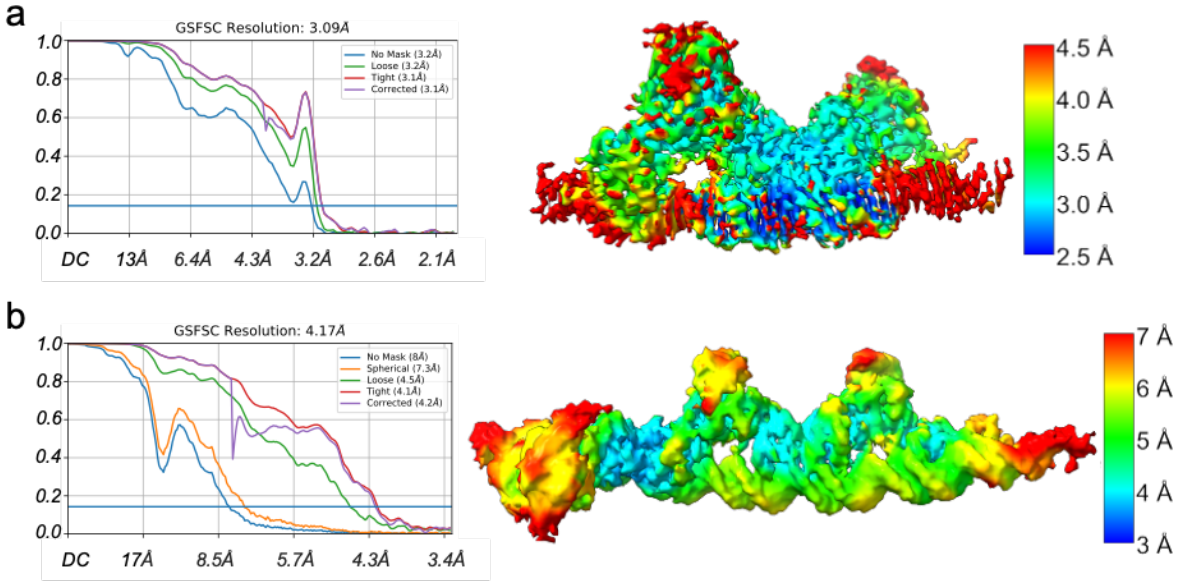


**Supplementary Figure 3. Resolution of EM density maps used in this study.** Gold-standard Fourier shell correlation (GSFSC) curve (left) and local resolution electron density maps (right) for (a) $(MntR_2)_2$-P84 and (b) $(MntR_2)_4$-P84 complexes showcasing that central region on the $(MntR_2)_2$-P84 complex has the highest resolution, about 2.5-3.0 Å, at the dimer-dimer and MntR-DNA interfaces at the center of the complex.

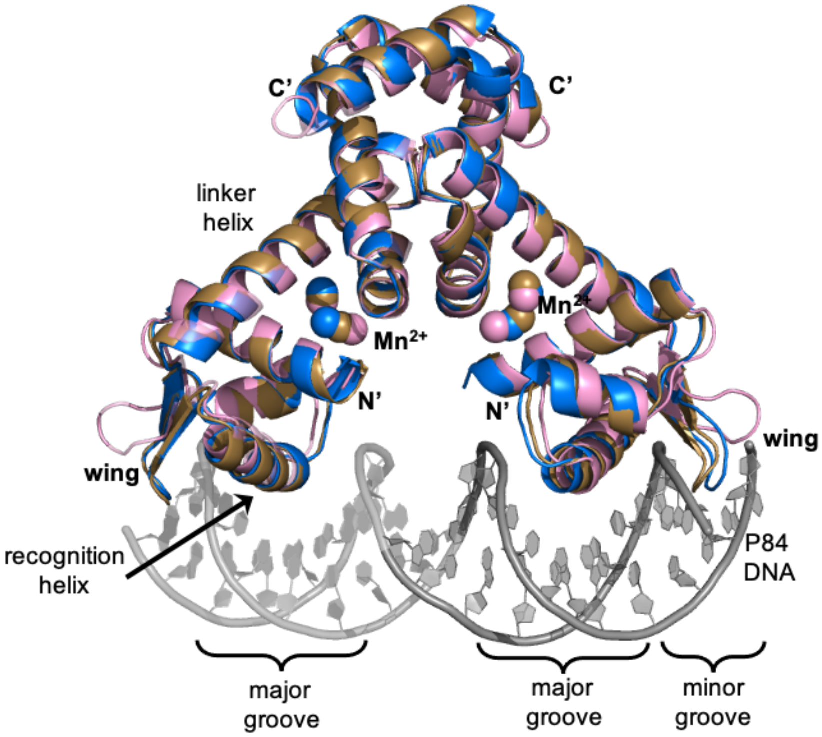


**Supplementary Figure 4. Structural alignments of MntR dimers with and without bound DNA.** Two MntR dimers from the $(MntR_2)_2$-P84 structure (PDB: 9C4C) were aligned to starting model of MntR dimer without the bound DNA[11] (PDB: 2F5C) in PyMOL[41] with RSMD values of 0.8 Å (tan) and 0.9 Å (blue).

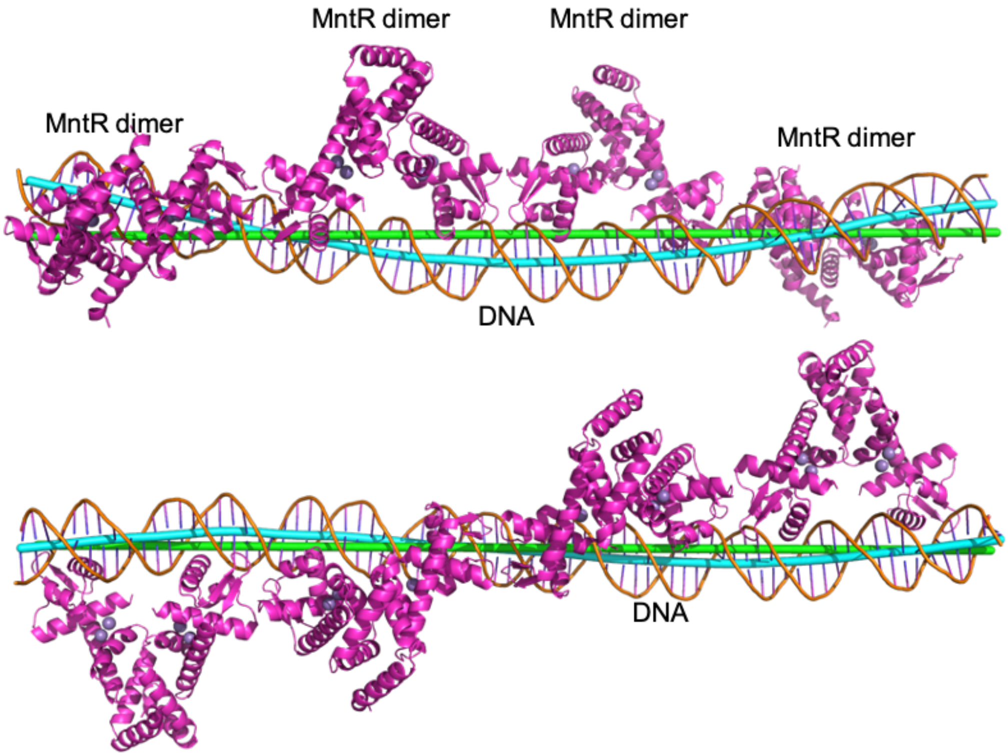


**Supplementary Figure 5. Two orthogonal views of the $(MntR_2)_4$-P84 complex with helix axes calculated by *Curves+* software.** The linear axes (shown as green tubes) are fit across all 78 modeled base pairs. The cyan-colored tube was calculated with changes in orientation at each base pair across the duplex. The DNA bends slightly toward the MntR dimers, with the most prominent inflection at points where MntR dimers interact.

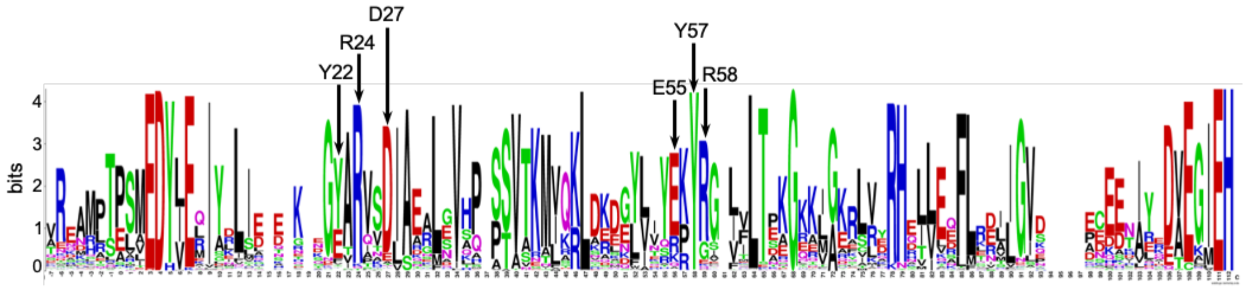


**Supplementary Figure 6. A Weblogo highlighting sequence conservation among MntR homologs.** Sequence analysis of MntR and its homologs was based on the results of a BlastP (Ref) search against the *Bs*MntR sequence using the refseq_select database, extending the results to an E value of 1e-15. The analysis resulted in 2334 sequences that were used to create a logo using Weblogo (http://weblogo.berkeley.edu/logo.cgi). Conservation level correlates with the diversity of residues at a specific position, with the symbol's height indicating the frequency of occurrence within the sequence. Some of the *B. subtilis* MntR amino acids of interest are highlighted with black arrows.

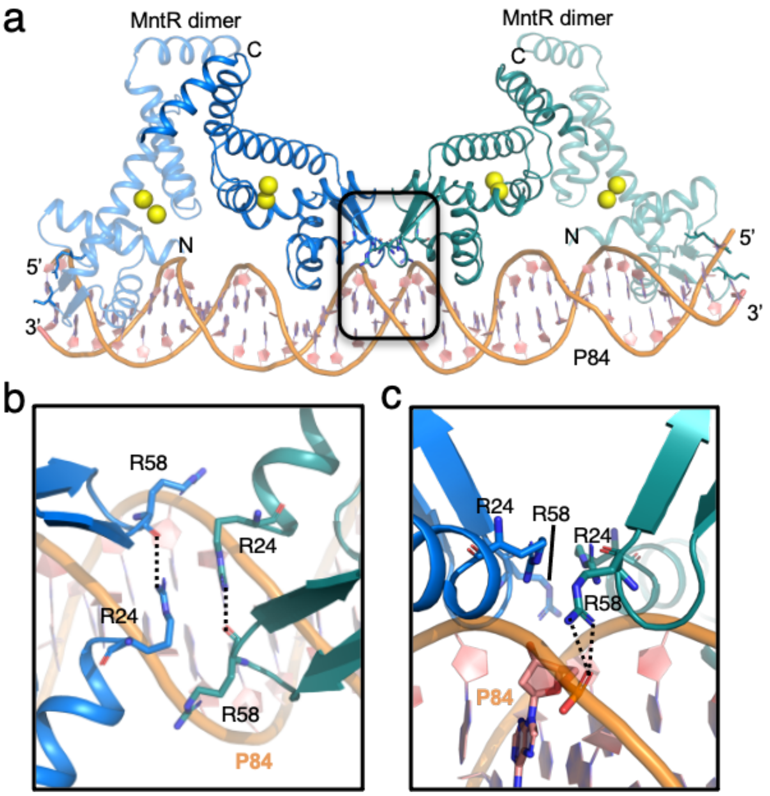


**Supplementary Figure 7. Arginine cluster at the MntR dimer-dimer interface.** (a) A cartoon representation of the cryo-EM structure of the $(MntR_2)_2$-P84 complex with two MntR dimers highlighted in green and blue, with each dimer bound to 4 $Mn^{2+}$ ions (yellow spheres). The MntR dimers are interacting with a portion of P84 (orange). Arg24 and Arg58 are highlighted in the cartoon. (b) Close up view from the top of the boxed region in (a) showcasing the network of arginine residues at the MntR interdimer interface. Arg24 is H-bonded to the backbone carbonyl of Arg58 on the same MntR subunit and Arg24 on the two adjacent MntR subunits are stacked against each other. (C) An alternate view of the (b) after a 90° rotation highlighting the H-bond (black dotted line) between Arg58 and the backbone phosphate group on P84. Note that all the four arginine residues form H-bonds with the backbone phosphate group on P84.

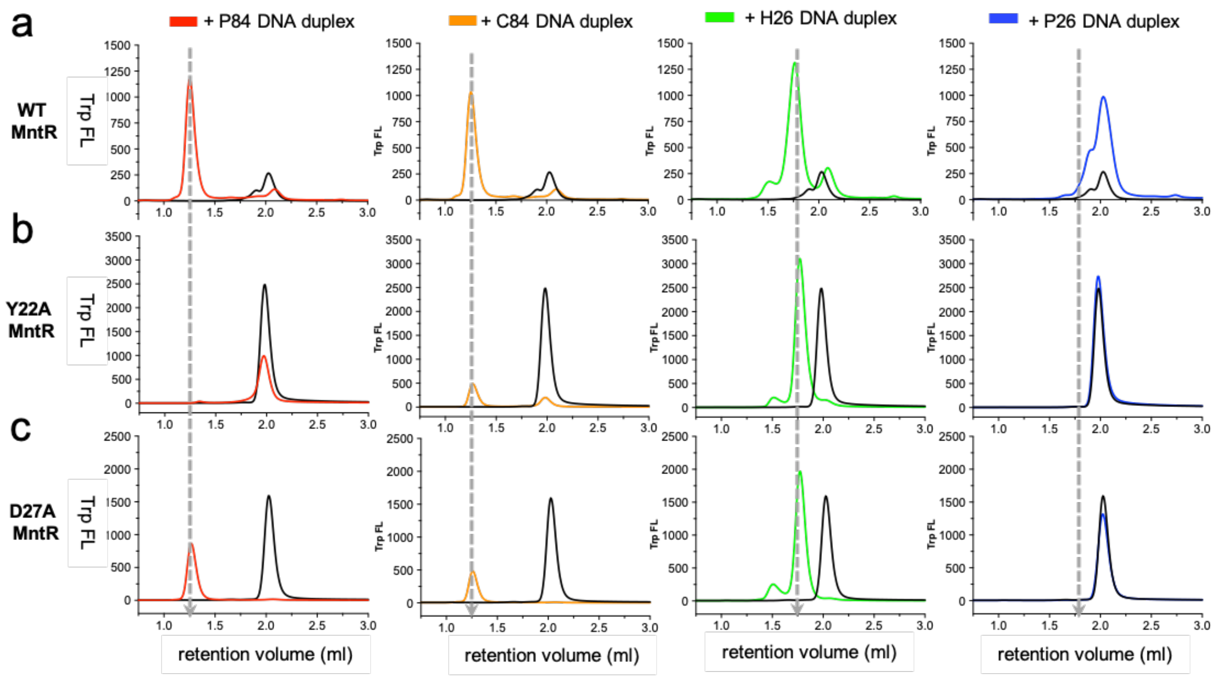


**Supplementary Figure. 8. Tryptophan FSEC experiments** were used to study binding of (a) WT MntR, (b) Y22A MntR and (c) D27A MntR with P84 (red), C84 (orange), H26 (green) and P26 (blue). The black traces represent the chromatograms of MntR proteins alone. The colored traces represent the chromatogram of MntR variants in the presence of the DNA duplex indicated. The broken grey lines are used to mark the retention volume of the MntR-DNA complex.
